# Ephrin receptor A4 (EphA4) is a new Kaposi’s sarcoma-associated herpesvirus virus entry receptor

**DOI:** 10.1101/510651

**Authors:** Jia Chen, Xianming Zhang, Samantha Schaller, Theodore S. Jardetzky, Richard Longnecker

## Abstract

Kaposi’s sarcoma-associated herpesvirus (KSHV) is a human γ-herpesvirus associated with the development of Kaposi’s sarcoma (KS). KSHV target cells include endothelial cells, B cells, monocytes, epithelial cells, dendritic cells, macrophages, and fibroblasts. KSHV entry into target cells is a complex multistep process and is initiated by the binding and interaction of viral envelope glycoproteins with the cellular receptors. In our current studies, we have found that EphA4 promotes KSHV gH/gL-mediated fusion and infection better than EphA2 in HEK293T cells indicating that EphA4 is a new KSHV entry receptor. To confirm that epithelial cells express EphA2 and EphA4, we analyzed the expression of EphA2 and EphA4 in epithelial cells, endothelial cells, B cells, monocytes, fibroblasts using RNA-seq data analysis of existing data sets. We found these cell types broadly express both EphA2 and EphA4 with the exception of monocytes and B cells. To confirm EphA4 is important for KSHV fusion and infection, we generated EphA2 and EphA4 single and double knockout cells. We found that both EphA2 and EphA4 play a role in KSHV fusion and infection, since EphA2/EphA4 double knockout cells had the greatest decrease in fusion activity and infection compared to single knockout cells. Fusion and infection of KSHV was rescued in the EphA2/EphA4 double knock cells upon overexpression of EphA2 and/or EphA4. EphA2 binds to both EBV and KSHV gH/gL; however, EphA4 binds only to KSHV gH/gL. Taken together, our results identify EphA4 as a new entry receptor for KSHV.

**Importance:** The overall entry mechanism for herpesviruses is not completely known including that for the human γ-herpesviruses Kaposi’s sarcoma-associated herpesvirus (KSHV) and Epstein-Barr virus (EBV). To fully understand the herpesvirus entry process, functional receptors need to be identified. In our current study, we found that EphA4 can also function for a KSHV entry receptor along with EphA2. Interestingly, we found that EphA4 does not function as an entry receptor for EBV whereas EphA2 does. The discovery of EphA4 as KSHV entry receptor has important implications for KSHV pathogenesis in humans and may prove useful in understanding the unique pathogenesis of KSHV infection in humans and may uncover new potential targets that can be used for the development of novel interventional strategies.

## Introduction

Herpesviruses are enveloped double-stranded DNA viruses capable of infecting a wide range of hosts and cause a variety of diseases (1). There are nine human herpesviruses that infect humans establishing lifelong latent infections (2). The γ-herpesviruses Kaposi’s sarcoma-associated herpesvirus (KSHV) and Epstein-Barr virus (EBV) are associated with human cancer (1, 3). Kaposi’s sarcoma (KS) is a cancer that develops from the endothelium that lines lymph or blood vessels. It usually appears as tumors on the skin or on mucosal surfaces such as inside the mouth, lungs, liver, and gastrointestinal tract. Skin lesions usually cause no symptoms; however, KS may become life-threatening if the lesions develop in essential organs such as the lungs, liver, or digestive tract (3).

KSHV infection is essential for the development of KS. The first step of KSHV infection is entry into target cells which is a complex multistep process. KSHV target cells in humans include endothelial cells, B cells, monocytes, epithelial cells, dendritic cells, macrophages, and fibroblasts (4). Entry of herpesviruses into target cells is initiated by the binding and interaction of viral envelope glycoproteins with cellular receptors, leading to either fusion of the viral envelope with the host cell membrane or endocytosis of viral particles and subsequent fusion of the viral envelope with an endocytic membrane for capsid release (5). Understanding the virus entry process may aide in the development of novel entry inhibitors and vaccines. Multiple KSHV receptors have been identified including integrins (α3β1, αVβ3, αVβ5), xCT (cystine/glutamate transporter), intracellular adhesion molecule-3 (ICAM-3), dendritic cell-specific intercellular adhesion molecule-grabbing non-integrin (DC-SIGN), and EphA2 (6–10). The integrin receptors and xCT are likely non-essential for infection and only required for the initial binding of the virus to cells (11), whereas EphA2 is essential because it triggers fusion upon virus binding to epithelial cells (10). The role of EphA2 in EBV entry was only recently identified by us and another group (12, 13). Both EBV and KSHV are associated with epithelial cell cancers, indicating that the engagement of EphA receptors could be a key commonality in the development of these malignancies.

In our current studies, we found that EphA4 functions as an entry receptor for KSHV. It is intriguing that the two human γ-herpesviruses, EBV and KSHV, use EphA proteins as entry receptors. Both EphA2 and EphA4 belong to the Eph receptor family, a large family of receptor tyrosine kinases (RTKs). The Eph receptor family contains 14 members and is divided into two classes, A and B, based upon sequence similarity and affinity with the 9 ephrin ligands (14). The Eph receptors and their ligands have bi-directional signaling capacity, indicating that they can serve as both receptors and ligands (14). The Ephrin receptors transduce signals from the cell exterior to the cell interior by ligand-induced activation of their kinase domain (15). The function of the Eph family includes boundary formation, cell migration, axon guidance, synapse formation, angiogenesis, proliferation, and cell differentiation (15, 16). Eph receptors have been implicated in regulating cell migration, adhesion, proliferation, and differentiation (16). Altered expression patterns of Eph receptors and ephrins (Eph receptor ligands) have been correlated with tumor behavior, such as invasiveness, vascularization, metastatic potential, and patient prognosis (17). Overexpression of EphA2 and EphA4 has been reported in gastric cancer, breast cancer, colon cancer, and prostate cancer (17–24). In our current studies, we investigated and found that EphA4 functions as an entry receptor of KSHV. Overall, our current findings broaden the knowledge of KSHV entry process and herpesvirus entry in general and may facilitate the development of potential entry inhibitors targeting KSHV infection.

## Results

### EphA4 promotes both KSHV cell-cell fusion activity and infection

We recently discovered that EBV uses EphA2 but not EphA4 as an epithelial cell entry receptor (12). EphA2 had previously been shown to function as a KSHV entry receptor for endothelial and epithelial cells (10). In our current studies, we investigated if EphA4 may also function in KSHV entry. Key in these studies was the use of EBV gB in the KSHV fusion assay in place of KSHV gB. Our previous work had shown that the KSHV cell-cell fusion assay was not as robust as the EBV fusion assay, making it difficult to obtain reproducible and significant data (25, 26). By replacing KSHV gB with EBV gB, KSHV fusion is greatly enhanced with fusion levels much higher than that mediated by the EBV glycoproteins (26). The exact nature of this enhancement is unknown, but likely is due to differences in the fusion activity of EBV gB compared to KSHV gB although a role of gH/gL can not be excluded.

To determine if EphA4 may trigger fusion, we transfected EBV gB with KSHV gH/gL and EBV gH/gL as a control in CHO-K1 cells and quantified fusion with HEK293T target cells overexpressing either EphA2 and EphA4. We found that overexpression of EphA4 induced higher fusion activity for the EBV gB and KSHV gH/gL combination compared to when EphA2 was tested (Fig. 1A). Our previous data showed that EphA2 but not EphA4 is required for EBV fusion activity (12) similar to the results shown when EBV gB and EBV gH/L are used (Fig 1A). This published data combined with the results shown in Fig. 1A suggest a specificity of EphA4 for KSHV gH/gL for fusion function when compared to EBV gH/gL. To further examine if EphA4 is sufficient to induce fusion for KSHV gH/gL, we examined fusion activity using a split GFP fusion assay and readily detected fusion as monitored by the appearance of green cells indicative of fusion activity only when KSHV gH/gL and EBV gB expressing cells were overlaid with cells that overexpress EphA2 or EphA4 (Fig. 1B). Similarly, EphA4 induced more fusion activity compared to EphA2, consistent with our observation of greater fusion in HEK293T cells transfected with EphA4. To examine the effect of EphA4 on KSHV infection of epithelial cells, we transfected HEK293T cells with control plasmid, EphA2, EphA4, and EphA2/EphA4 and then infected the cells with KSHV virus expressing GFP. Flow cytometry showed increased infection of KSHV in the presence of either EphA2 or EphA4 and higher levels of fusion when both EphA2 and EphA4 were transfected. (Fig. 1C). We also examined infection by fluorescence microscopy (Fig. 1D) which was consistent with the flow cytometry data (Fig. 1C).

**Figure 1.**
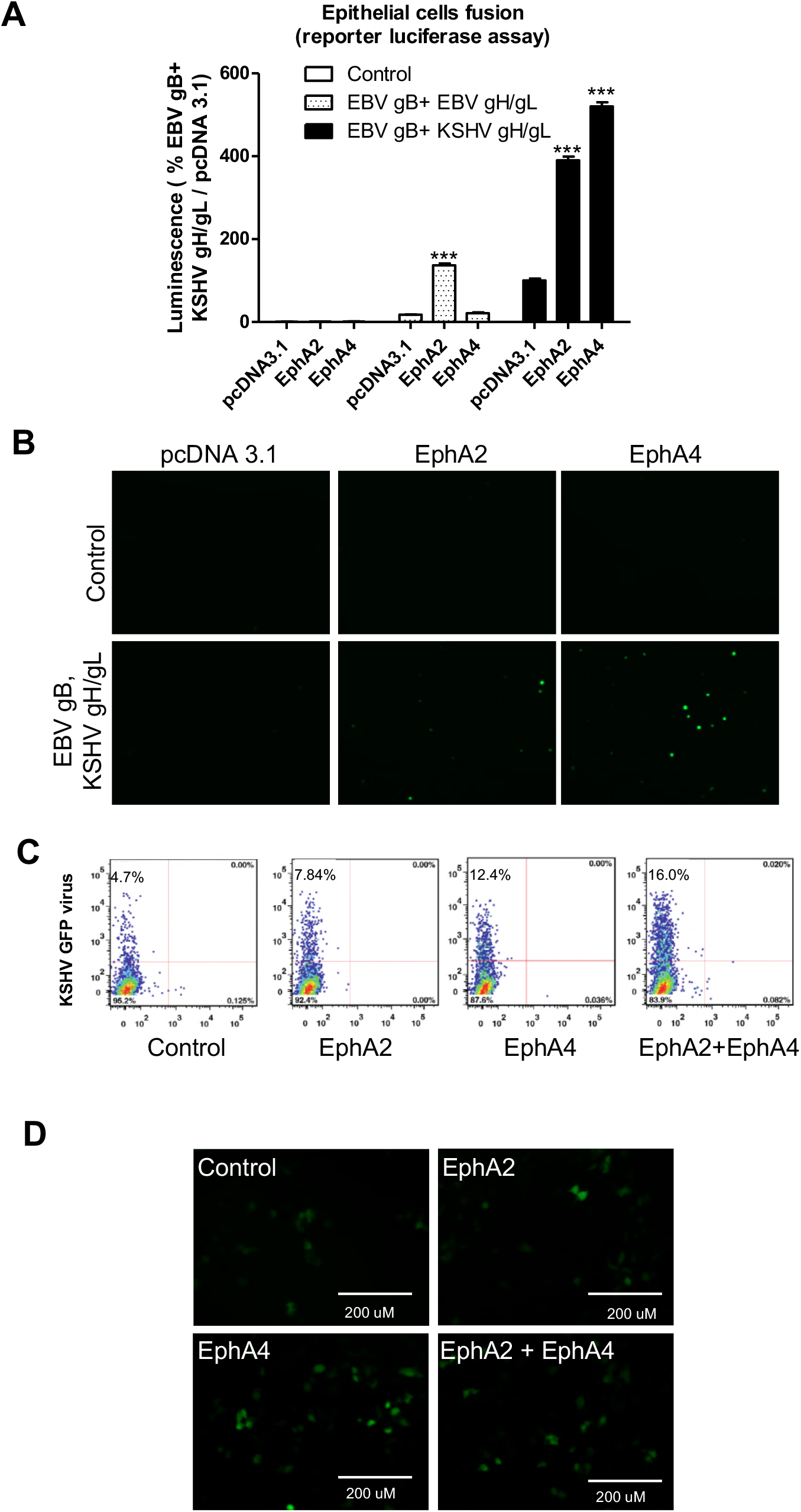
EphA4 promotes both KSHV infection and virus-free cell-cell fusion. **A**, CHO-K1 cells were transfected with T7 luciferase plasmid and either control plasmid, or EBV gH/gL, EBV gB, or KSHV gH/gL. Transfected CHO-K1 cells were overlaid with HEK293T cells transfected with pcDNA3.1, EphA2 or EphA4 together with T7 polymerase. Fusion activity was standardized to EBV gB, KSHV gH/gL fusion with HEK293T cells transfected with control pcDNA3.1 which was set to 100%. ***P<0.001 (ANOVA followed by post-hoc Tukey’s multiple comparison test), compared to pcDNA 3.1. **B**. 2.5 x 10^5^ CHO-K1 cells transfected with Rluc8 1-7 luciferase plasmid together with either control plasmid, EBV gH/gL, EBV gB, or KSHV gH/gL, EBV gB, were overlaid with 2.5 x 10^5^ CHO-K1 cells transfected with pcDNA3.1, EphA2, or EphA4 together with Rluc8 8-11. Green cells, indicative of fusion, were visualized and captured with an EVOS fluorescence microscope. **C**, HEK293T cells were transfected with pcDNA3.1, EphA2, or EphA4. 24 hours post-transfection, 5×10^4^ cells were seeded into a 48-well plate. 24 hours later, the cells were infected with concentrated KSHV. After an additional 24 hours, the infected cells were analyzed by flow cytometry (C) or visualized by microscopy and images captured with an EVOS fluorescence microscope (**D**).

### EphA2 and EphA4 are expressed in various KSHV target cells and both function in KSHV entry

KSHV has broad tropism since its genome and transcripts can be detected in vivo and in vitro in a variety of cell types (27). To confirm that EphA4 is expressed in cells infected by KSHV, we analyzed existing RNA-seq data sets from B cells, monocytes, epithelial cells, fibroblasts, and endothelial cells available from SRA database (https://www.ncbi.nlm.nih.gov/sra). Neither EphA2 nor EphA4 was expressed abundantly in monocytes, indicating that entry of KSHV into monocytes may use other receptors (Fig. 2 A-D). Whereas, EphA2 and EphA4 were expressed in B cells, epithelial cells, fibroblasts, and endothelial cells (https://www.proteinatlas.org/ENSG00000116106-EPHA4/tissue), consistent with KSHV using EphA2 and EphA4 as primary entry receptors in these cell types. To further confirm that EphA4 can serve as a cellular receptor for KSHV infection, we generated EphA2 and EphA4 single and double knockout cells using the CRISPR/Cas9 system in HEK293T cells. Following knockout, EphA2 cell surface expression was determined by flow cytometry. As expected, there was a lack of EphA2 expression as analyzed by flow cytometry in the EphA2 single knockout cells and in the EphA2/EphA4 double knockout cells but not in the EphA4 knockout cells and WT cells (Fig. 3A). We analyzed EphA4 expression by Western blotting since the available antibodies did not work well for flow cytometry. EphA4 expression was not detected in EphA4 single knockout cells and in the EphA2/EphA4 double knockout cells (Fig. 3B). We next examined the effect of EphA2 and EphA4 knockout on KSHV fusion. We found that knockout of EphA2 and EphA4 individually dramatically decreased fusion activity (Fig. 3C). In the EphA2 and EphA4 double knockout cells, fusion activity was further decreased compared to single knockout cells (Fig. 3C). When EphA2 or EphA4 were overexpressed in the double knockout cells, fusion activity was rescued (Fig. 3D). These data confirmed that both EphA2 and EphA4 are functional for KSHV fusion. Finally, we investigated if EphA2 and EphA4 expression restored KSHV infection in the double knockout cells. When EphA2 and EphA4 were individually transfected into the double knockout cells, infection with KSHV was partially rescued when compared to levels observed in HEK293T cells (Fig. 3E). The level of infection in EphA2 expressing cells was just above background levels in contrast to the EphA4 in which the level of infection was higher (Fig. 3E). Overall, the fusion and infection results presented in Figure 3 indicate that both EphA2 and EphA4 both function as receptors with EphA4 being the better receptor in the assays used in our current studies.

**Figure 2.**
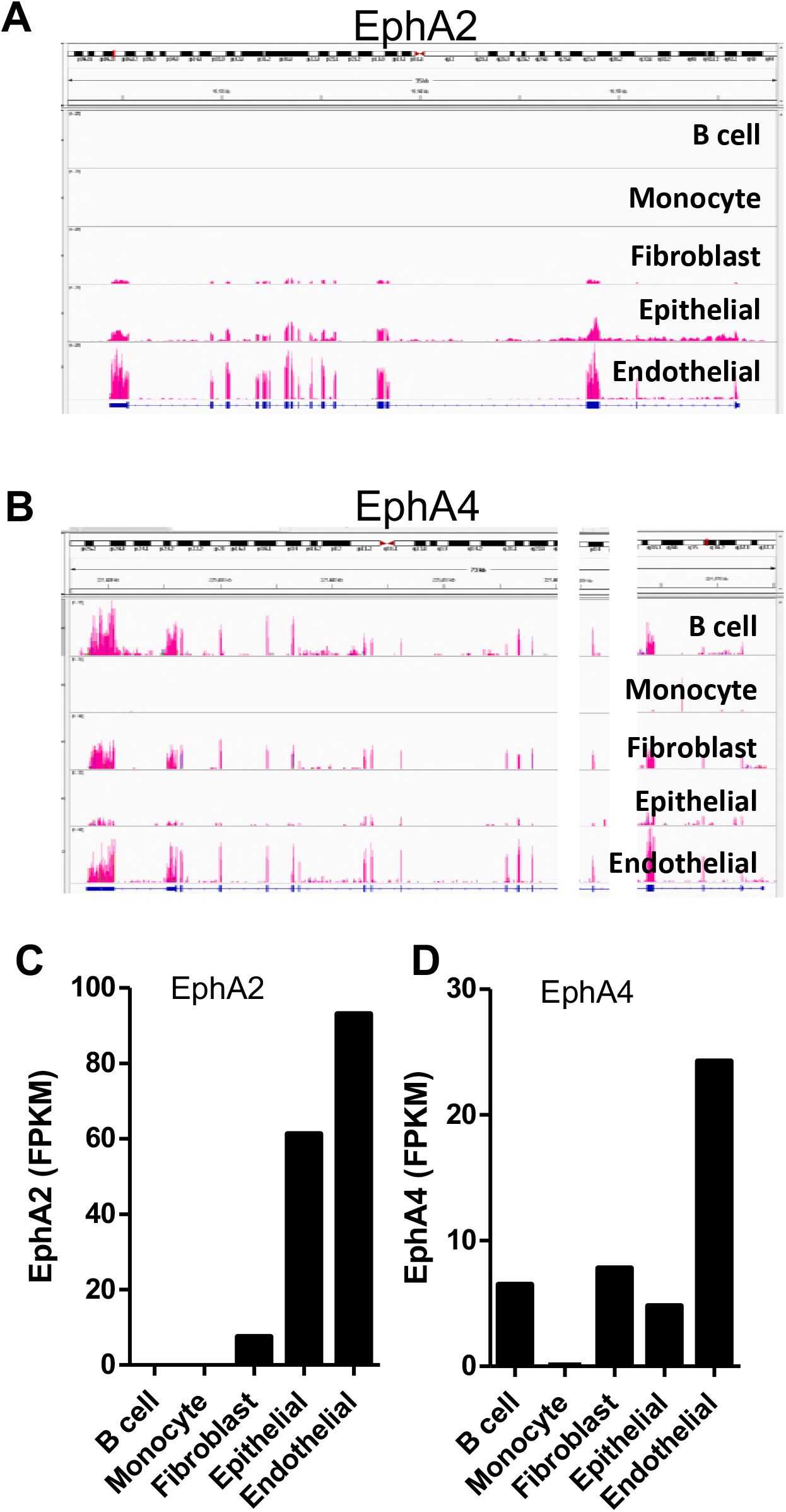
EphA2 and EphA4 expression in KSHV target cells. **A** and **B** The distribution of EphA2 (**A**) and EphA4 (**B**) sequencing reads across EphA2 or EphA4 exons. BAM formatted files were generated by alignment of RNA-seq data from various cell types infected by KSHV. RNA-seq data were obtained from Sequence Read Archive database (NIH) and aligned data was then loaded into the Integrative Genomics Viewer (Broad Institute) to acquire the transcript map with exon reads shown in red. Two chromosomal regions, in which no reads for EphA4 were detected, were removed and are shown as white bars allowing the transcript map to fit within the figure. **C** and **D**, The mean FPKM (Fragments Per Kilobase of transcript per Million mapped reads) of EphA2 (**C**) and EphA4 (**D**) in various KSHV target cells. FPKM was determined using Cuffdiff software (Broad Institute).

**Figure 3.**
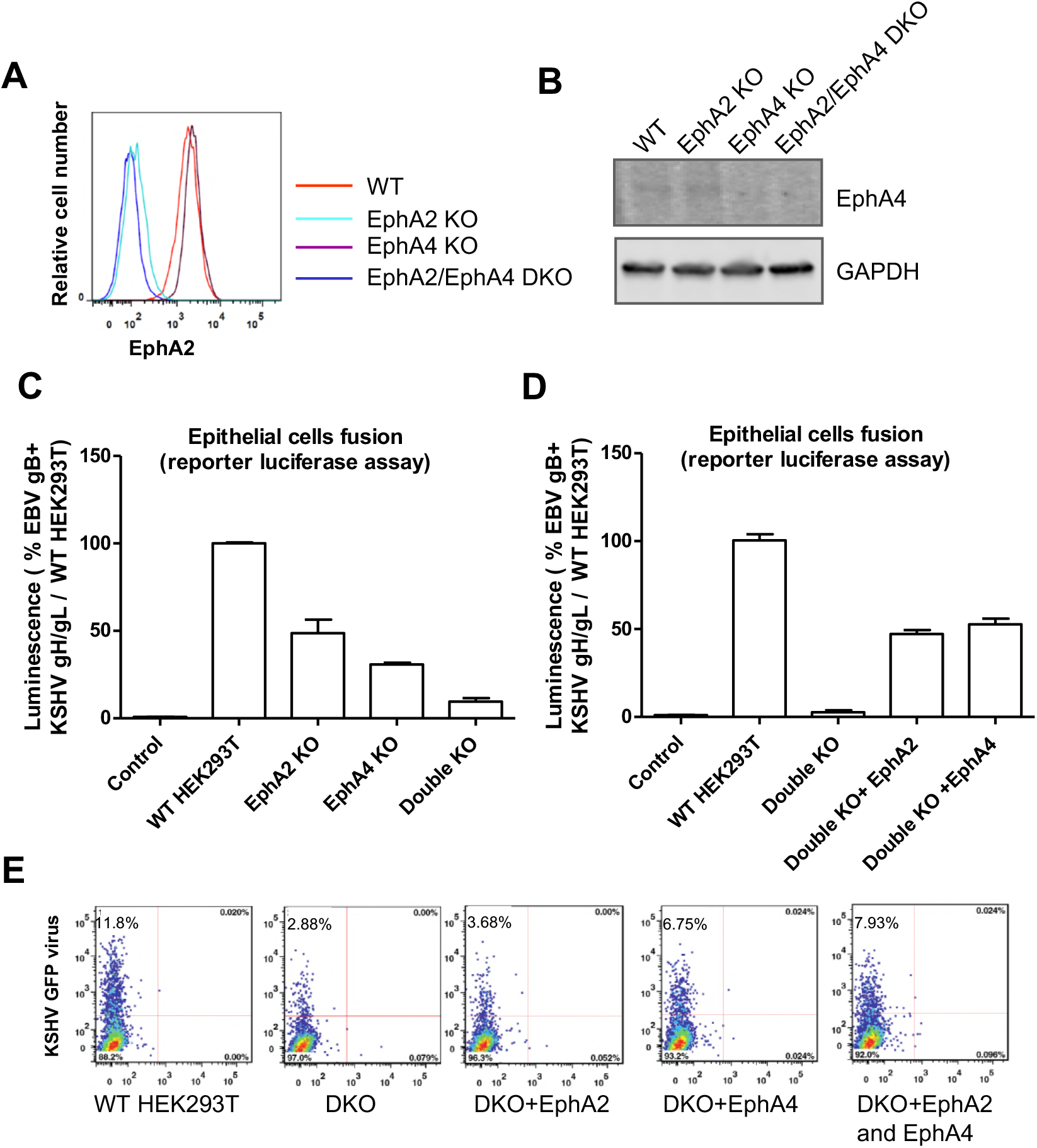
EphA4 is the potential epithelial cell receptor for KSHV. **A.** EphA2 cell surface expression in EphA2 and EphA4 single or double knockout (DKO) HEK293T cells by flow cytometry. The X axis represents the relative number of cells analyzed by flow cytometry with a particular level of EphA2 expression. The y-axis represents the level of expression within the analyzed cell population on a log scale. **B**, EphA4 expression in EphA2 and EphA4 single or double knockout HEK293T cells by Western blotting. GAPDH was used as loading control. **C**, For fusion function of knockout cell lines, CHO-K1 cells transfected with T7 luciferase and either a control plasmid or KSHV gH/gL, EBV gB were overlaid with EphA2 and EphA4 single or double knockout cells (**C**), or EphA2/A4 double knockout cells overexpressing EphA2, EphA4 or EphA2/EphA4 (**D**), that were transfected with T7 polymerase. **E**, WT or EphA2 and EphA4 double knockout cells were transfected with EphA2 or EphA4, and infected with KSHV as described in the material and methods. 24 hours later, cells were analyzed by flow cytometry for GFP expression to identify infected cells (**E**).

### Both EphA2 and EphA4 bind to KSHV gH/gL

Previous studies indicated that both EphA2 and EphA4 can bind KSHV gH/gL by co-immunoprecipitation, but the function of EphA4 to mediate entry was not studied (28). To confirm the previously observed binding of KSHV gH/gL with EphA2 and EphA4, we used three different methods. We first transfected CHO-K1 cells with control plasmid, EBV gH/gL, or KSHV gH/gL. Twenty-four hours later, the cells were detached and seeded in a 96-well plate in triplicate or in a 6-well plate. Supernatants from cells transfected with the soluble forms of EphA4-Fc and EphA2-Fc were added to cells transfected with KSHV or EBV gH/gL at 4° C. The binding of EphA2 or EphA4 with EBV or KSHV gH/gL was then determined by CELISA or western blotting (Fig. 4A and 4B). The CELISA data showed that EphA2 can bind to both EBV and KSHV gH/gL, but with higher levels for KSHV gH/gL (Fig. 4A). However, EphA4 only bound to KSHV gH/gL and not EBV gH/gL (Fig. 4A). This is consistent with the observation that EphA4 expression does not increase EBV fusion. When soluble EphA2-Fc is co-expressed with EBV gH/gL, KSHV gH/gL, or control vector transfected cells, soluble EphA2-Fc can bind to both EBV and KSHV gH/gL as detected by CELISA (Fig. 4C). However, soluble EphA4-Fc can only be detected when KSHV gH/gL are co-expressed and not with EBV gH/gL (Fig. 4D). These data confirmed that KSHV gH/gL can bind to both EphA2 and EphA4, whereas EBV gH/gL binds better to EpHA2 when compared to EphA4 (Fig. 4A). There was some difference in the efficiency of EphA2 binding to KSHV gH/gL and EBV gH/gL when Fig. 4A and 4C are compared. This is likely a result of the different methods used to detect binding as described in the legend for Fig. 4 and the Materials and Methods.

**Figure 4.**
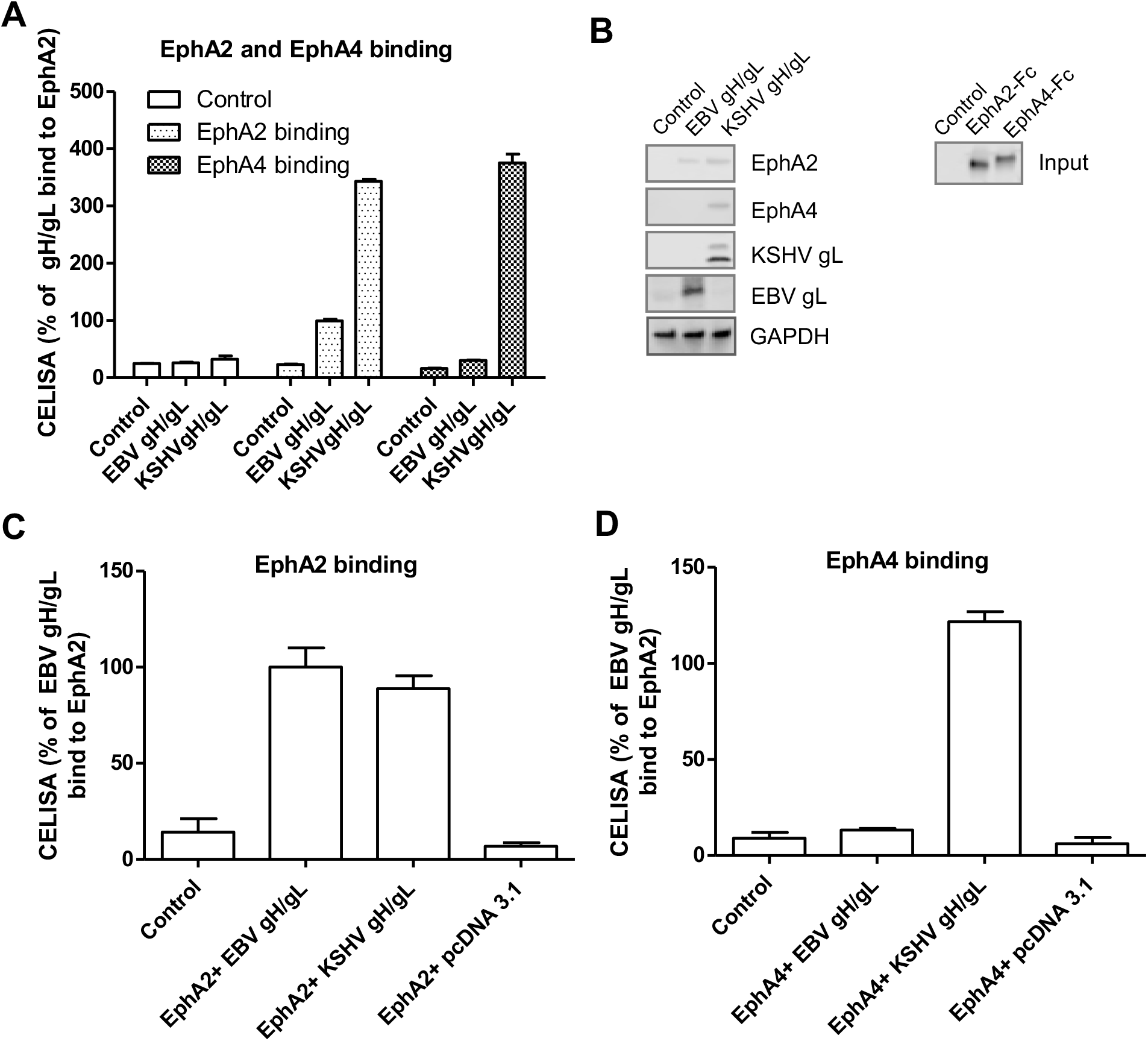
Both EphA2 and EphA4 bind to KSHV gH/gL. (**A**) CHO-K1 cells were transiently transfected with control pcDNA3.1, EBV gH/gL, or KSHV gH/gL plasmids. Soluble EphA2-Fc or EphA4-Fc was prepared by transfecting CHO-K1 cells with EphA2-Fc or EphA4-Fc plasmid constructs and media supernatants containing EphA2-Fc or EphA4-Fc were overlaid on HEK293T cells expressing EBV gH/gL or KSHV gH/gL in 96-well plates in triplicate for 2 hours at 4°C and bound protein was detected using anti-human IgG which recognizes the Fc portion of the Eph2-Fc and EphA4-Fc by CELISA. **B**. CHO-K1 cells seeded in 6-well plates were transfected with control plasmid pcDNA3.1, EBV gH/gL, or KSHV gH/gL with gL containing a His-tagged as indicated. After 24 hours, cells were washed twice with ice-cold PBS and incubated with supernatants from pcDNA3.1 (control plasmid), EphA2-Fc, or EphA4-Fc transfected cells isolated 24 hours post-transfection for 2 hours at 4°C. The cells were then washed with ice-cold PBS three times and lysed with 200 ul of 1X SDS lysis buffer. Proteins bound to the cells expressing EBV gH/gL or KSHV gH/gL were then analyzed using antibodies to the Fc region of the EphA2 and EphA4 fusions by Western blotting. GAPDH was used as a loading control. Expression of the KSHV gH/L or EBV gH/gL complex was monitored by analyzing KSHV gL or EBV gL expression using anti-His antibodies directed against a His-tagged added to KSHV gL or polyclonal antibodies directed against EBV gL. **C and D**, CHO-K1 cells seeded in 6-well plates were transfected with control plasmid pcDNA3.1, EBV gH/gL, or KSHV gH/gL together with EphA2-Fc or EphA4-Fc. Transfected cells were seeded in 96-well plates in triplicate post-transfection. The cells were then washed with ice-cold PBS three times and gH/gL-associated EphA2 (**C**) or EphA4 (**D**) was determined by CELISA using anti-human IgG antibodies.

### The ectodomain of EphA2 and EphA4 are interchangeable for KSHV fusion activity and EphA4 kinase activity is not needed for KSHV fusion activity

EphA4 is a membrane protein with four different ectodomain regions including a ligand binding domain (LBD), a cysteine rich region (CYS), and two fibronectin regions (FBN). EphA4 and EphA2 share about 51% similarity at the amino acid level. The kinase domain is located within the cytoplasmic tail domain. Previous studies by Hahn et al indicated that the ectodomain of EphA2 is important for binding with KSHV (10). Our previous results with EBV found that the ligand binding domain (LBD) is important EBV fusion function. We demonstrated this by swapping the LBD of EphA2 and EphA4 to generate EphA2A4 or EphA4A2 LBD chimeras (12). We used the same constructs to investigate if the LBDs of EphA2 and EphA4 are interchangeable for KSHV gH/gL. Overall, all of the chimeras worked well in KSHV fusion (Fig. 5A and 5B). Interestingly, the EphA4A2 chimera functioned better in fusion than the EphA2A4 chimera indicating that the LBD domain may be responsible for the greater fusion activity observed for EphA4 when compared to EphA2.

**Figure 5.**
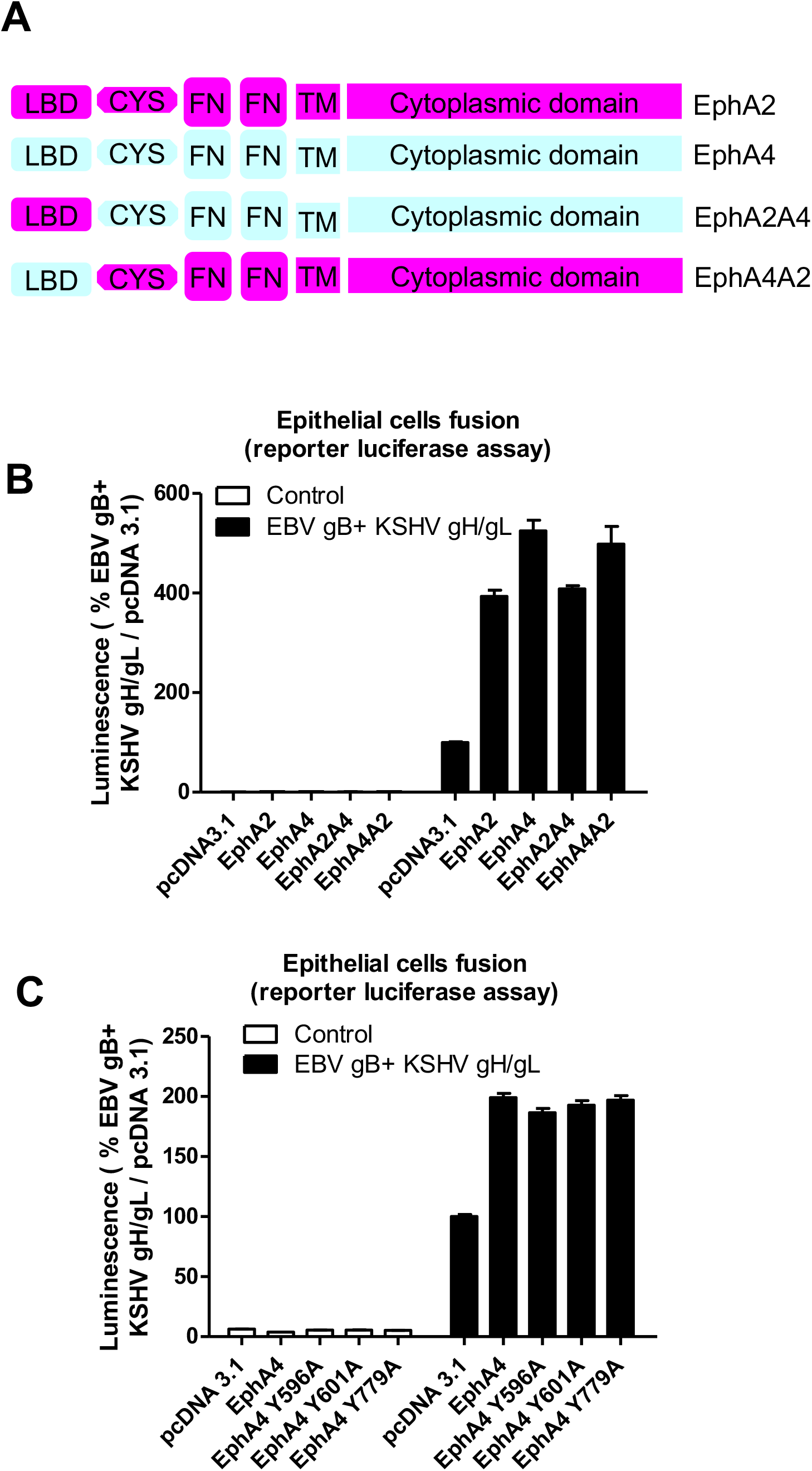
The ectodomain of EphA2 and EphA4 are interchangeable for KSHV fusion activity and the kinase activity of EphA4 is not needed for KSHV fusion activity. **A.** Schematic drawing of the EphA2, EphA4, and, EphA2/EphA4 chimeras including the ligand binding domain (LBD). **B.** KSHV fusion with HEK293T cells transfected with control plasmid pcDNA3.1, EphA2, EphA4 or EphA2 and EphA4 chimeras as indicated. **C**, KSHV fusion with HEK293T cells transfected with control plasmid pcDNA3.1, EphA4, or EphA4 kinase-dead mutants as indicated. Fusion activity of HEK293T cells transfected with pcDNA 3.1 was set to 100% (**B** and **C**).

Previous studies had demonstrated that the kinase region of EphA2 was important for KSHV endocytosis (10) and subsequent entry. Our previous results demonstrated the EphA2 kinase function was not important for EBV mediated cell-cell fusion (12). To investigate if the kinase function was also nonessential for KSHV cell-cell fusion, we constructed three EphA4 kinase-dead mutants based on previous studies (27, 28) similar to the EphA2 constructs that we used in our previous study with EBV (12). In these constructs, tyrosine 596, 602, and 779 were all mutated individually to alanine. These three mutants all lack kinase activity based on prior mutagenesis studies (29, 30). The three EphA4 kinase dead mutants were transfected into HEK293T cells. Fusion activity of the mutants compared to WT EphA4 (Fig. 5C) were similar indicating that the kinase activity is not important for the fusion function of EphA4. The lower fusion observed in the pcDNA3.1 transfected cells is a result of the existing EphA2 and EphA4 expression in these cells since the EphA2/EphA4 double knockout cells were not used for this experiment.

## Discussion

KSHV infection is essential for the development of KS (3). This study has shown, similar to EphA2, EphA4 is a receptor that can directly interact with KSHV gH/gL. Moreover, we found that EphA4 functions better when compared to EphA2 since overexpression of EphA4 enhances KSHV fusion by approximately 33% and KSHV infection by 163% compared to EphA2 (Fig. 1). Both EphA2 and EphA4 are expressed in many KSHV target cells including, epithelial cells, fibroblasts, and endothelial cells (Fig. 2). Interestingly, EphA4 but not EphA2 is expressed in B cells, suggesting EphA4 might play a key role in KSHV B cell infection exclusively compared to EphA2. Knockout of both EphA2 and EphA4 greatly decreased fusion and infection of KSHV, while overexpression of EphA2 and EphA4 alone or together rescued the fusion and infection of EphA2 and EphA4 double knock out cells (Fig. 3). These findings indicate that EphA4 might be a novel factor important for KSHV infection and may play a role in KS pathogenesis.

Recently, EphA2 has been identified as the receptor for several viruses and as an epithelial cell pattern recognition receptor for fungal B-glucans (31). The human blood-brain barrier internalizes Cryptococcus neoformans via EphA2 (32). EphA2 is a receptor for KSHV and EBV and a cellular co-factor for hepatitis C virus entry (33). Additionally, ephrinB2, one ligand of this family, has been identified as a receptor for Nipah virus (34, 35). Thus, the Eph family and its ligands are entry factors for many pathogens.

Cell entry by KSHV is a multistep process involving viral envelope glycoproteins as well as cellular receptors and other cofactors leading to the merger of virus and host membranes (5). The binding receptors for KSHV include surface heparan sulfate (HS), which promotes a charge-based interaction between virus glycoproteins and the cell surface. Integrins such as α3β1, αVβ3, and αVβ5 play a crucial role in KSHV infection (6). xCT, a 12-transmembrane glutamate/cystine exchange transporter protein, and DC-SIGN have also been reported to be entry receptors for KSHV (7–9). More recently, EphA2 was identified as receptor that mediates KSHV infection of epithelial and endothelial cells (10). After the identification of EphA2 as the receptor for KSHV, Hahn, et. al screened the interaction of 14 Eph proteins with KSHV gH/gL. They found that other Eph family members also interact with KSHV gH/gL, including EphA4, EphA5, and EphB8 (28).

Like EBV gH/gL and HSV gH/gL, KSHV gH/gL forms a non-covalently linked complex (36). The function of gH/gL is to provide a key function in herpesvirus fusion to trigger gB activation and subsequent membrane fusion. Our previous results found that compared to EBV gB, KSHV gB is a poor fusogen (26). In our current study, we used this enhanced fusion function of EBV gB with KSHV gH/gL to increase fusion activity to facilitate our current studies. We found that while only EphA2 can enhance the fusion activity of EBV gH/gL, both EphA2 and EphA4 can enhance the fusion activity of KSHV gH/gL (Fig. 1A), indicating the specificity of EphA4 for KSHV gH/gL. Using a split GFP assay, we found that overexpression of EphA2 or EphA4 can induce fusion compared to control cells, indicating that EphA2 or EphA4 alone is sufficient to serve as the receptor for KSHV gH/gL (Fig. 1B). In WT HEK293T cells, overexpression of either EphA2 or EphA4 or both can also enhance KSHV infection (Fig. 1C, 1D).

EphA4 is widely expressed in different tissues and cell types (3). Using RNA-seq analysis we found that EphA4 is expressed in B cells, fibroblasts, epithelial cells, and endothelial cells (Fig. 2) suggesting a role for EphA4 in the infection of epithelial cells. To confirm this, we generated EphA2 and EphA4 single and double knockout cells. Fusion activity was decreased in both EphA2 (approximate 50% decrease) and EphA4 single knockout cells (approximate 70% decrease) (Fig. 3C). Fusion activity was drastically decreased in the EphA2 and EphA4 double knockout cells (90% decrease) (Fig. 3C). Transfection of the double knockout cells with EphA2 or EphA4, and especially both EphA2 and EphA4, rescued infection (Fig. 3E and 3F). These results together indicate that both EphA2 and EphA4 play a role in KSHV fusion. A recent paper published by TerBush et al. reveals an integrin-independent route of KSHV infection and also suggests that multiple Eph receptors besides EphA2 can promote and regulate infection, consistent with our findings (11).

As a KSHV receptor, EphA2 binds to KSHV gH/gL at nM level (12, 37). Previous co-IP data also showed that KSHV gH/gL can bind to EphA2, EphA4, EphA5 and EphB8 (28). We confirmed that EphA4 can bind to KSHV gH/gL. Using three different methods, we showed that EphA2 can bind to both EBV gH/gL and KSHV gH/gL independent of gB (Fig. 4) and that only EphA4 can bind to KSHV gH/gL as previously shown (12, 28). Recently, Großkopf et al. found a conserved motif ELEFN within gH of KSHV and rhesus monkey rhadinovirus (RRV) which is important for the EphA2 and EphB3 binding (38). Mutation of the ELEFN motif in RRV and KSHV is sufficient for detargeting of KSHV from Eph family receptors and reduces infection of susceptible cells (38). The corresponding motif in EBV gH is DIEGH. Interestingly, the first three amino acids in this motif, DIE, have similar structural and biochemical characteristics to the ELE found in KSHV gH. In contrast the last two amino acids in the ELEFN, F and N are structurally and biochemically quite different than the G and H found in EBV gH suggesting that these two regions may be critical for KSHV to use EphA2 and EphA4 as receptors.

Our results showed that EphA4 promotes KSHV fusion approximately 33% better than EphA2 (Fig. 1A and Fig. 5B) for KSHV fusion. Both KSHV gH/gL and EBV gH/gL binds to the EphA2 LBD domain, and the interaction can be competed with Ephrins (12, 13, 37). In the current study we tested EphA2 and EphA4 LBD chimeras and found that the LBD of EphA2 and EphA4 is interchangeable for KSHV fusion. Interestingly, it is the LBD that determines fusion activity since the chimera containing the EphA4 LBD (EphA4A2) behaves more like wt EphA4 and this is also true for EphA2A4 chimera (Fig. 5B)

Previously, we examined if the kinase activity of EphA2 is important for the fusion of EBV gH/gL and did not observe any loss of fusion function for the kinase-dead EphA2 mutants (12). In the current study, we examined the kinase activity of EphA4 in the context of KSHV gH/gL fusion. Similarly, we did not observe any changes between WT EphA4 and the EphA4 kinase-dead mutants for fusion function. Previous work indicated that binding of KSHV gH/gL to EphA2 triggered EphA2 phosphorylation and endocytosis (10). Since fusion levels are not altered in the EphA2 and EphA4 kinase dead mutants, it is likely that the kinase function is important for infection by functioning in endocytosis following virus binding (10).

Both EphA2 and EphA4 are overexpressed in numerous malignancies including gastric cancer, breast cancer, colon cancer, and prostate cancer (17–24). Since both EBV and KSHV infections are associated with multiple malignancies along with the observation that both EBV and KSHV use EphA family receptors, it will be of interest to determine if EphA2 and EphA4 might play a role in the development of KSHV and/or EBV associated cancers not only as entry receptors. The identification of EphA4 as a KSHV entry receptor provides new opportunities to understand the tissue tropism of KSHV and strategies to limit KSHV infection in the human host.

## Materials and Methods

### Cell culture

Chinese Hamster Ovary (CHO-K1) cells (ATCC CCL-61) were grown in Ham’s F-12 medium (Corning) containing 10% heat-inactivated fetal bovine serum (FBS) (Corning) and 1% penicillin-streptomycin (100 U penicillin/mL, 100 μg streptomycin/mL; Sigma). Human Embryonic Kidney 293 T (HEK293T) cells (ATCC CRL-3216) or HEK293-T14 cells derived from HEK293T stably expressing T7 RNA polymerase (39) were grown in DMEM (Corning) with 100μg/mL zeocin (Invitrogen, for HEK293T cells expressing T7 RNA polymerase only), containing 10% heat-inactivated FBS and 1% penicillin-streptomycin, respectively. iSLK.219 KSHV cells (40) were kindly provided by Eva Gottwein and were grown in DMEM (Corning) containing 10% Tetracycline-free fetal bovine serum complex (Clontech) and 1% penicillin-streptomycin (100 U penicillin/mL, 100 μg streptomycin/mL; Sigma).

### Constructs

The EphA2 and EphA4 constructs (12) were a gift from Dr. Spiro Getsios (Northwestern University). The construction of the EphA2 and EphA4 LBD chimeras (EphA2A4 or EphA4A2) was previously described (12). Soluble EphA2-Fc and EphA4-Fc were cloned in a Fc construct a gift from Dr. Qing Fan (41). EphA2-Fc (His tag) constructs oligos EphA2-Fc F: GACTCGAGATGCAGGGCAAGGAAGTGGTACTG and EphA2-Fc R: GACTCGAGgtggtgatggtgatgatgGTTGCCAGATCCCTCCGGGGA. EphA4-Fc (His tag) constructs oligos EphA4-Fc F: GACTCGAGATGGCTGGGATTTTCTATTTC and EphA4-Fc R: GACTCGAGgtggtgatggtgatgatgTGTGGAGTTAGCCCCATCTCC.

His tagged KSHV gL was subcloned into the pSG5 vector using the following primers: KSHV gL EcoRI F: GCGAATTCCATGGGGATCTTTGCGCTATTT. KSHV His gL BglII R: TAAGATCTGTTTAGTGGTGATGGTGATGATGTTTTCCCTTTTGACCTGCGTG EphA2 sgRNA constructs oligos: 5-AAACGTGTGCGCTACTCGGAGCCTC-3 and 5-CACCGGAAGCGCGGCATGGAGCTCC-3 were annealed and ligated into a lentiGuide-Puro plasmid (Addgene, # 52963). EphA4 sgRNA constructs: oligos 5-AAACCACAGTACATTTTTGGCACAC-3 and 5-CACCGTGTGCCAAAAATGTACTGTG-3 were annealed and ligated into a lentiGuide-Puro plasmid (Addgene, # 52963). Sequencing was performed for all constructs to confirm the correct sequence.

#### RNA-seq data assay

The RNA-seq data assay was performed as described in our previous paper (12). Briefly, the Sequence Read Archive (SRA) data of RNA-seq for B cells (SRR5048157 and SRR5048162), monocytes (SRR5048180 and SRR5048177), fibroblasts (SRR3192540 and SRR3192539), epithelial cells (SRR3192374 and SRR3192375), and endothelial cells (SRR3192370 and SRR3192369) were downloaded from the SRA database (https://www.ncbi.nlm.nih.gov/sra). The SRA data were transformed into the original FASTQ documents using NCBI SRA Toolkit fastq-dump. The original documents were further trimmed using FASTX and aligned to the reference genome using TopHat2. The differential expression analysis was performed using Cuffdiff software.

### Generation of EphA2 and EphA4 single and double KO cells

For EphA2 and EphA4 single and double KO cells, Cas9-expressing stable HEK293-T14 cells were established by infecting with lentivirus containing Cas9 for 24 hours. 24 hours later, the cells were changed to fresh medium with 5 ug/mL blasticidin for selection. After one week, single colonies were picked as previously described and expanded for 2-3 weeks (12). The Cas9 expression in these single cell colonies was analyzed using Western blotting using the Flag tag fused to Cas9. 2.5 x 10 ^5^ HEK293-T14-Cas9 cells per well in a 12-well plate were infected with lentivirus including control sgRNA, EphA2 sgRNA, or EphA4 sgRNA, either individually or combined together. The cells were selected with 2 ug/mL puromycin and coloned as single cells as previously described (12). After 2-3 weeks of expansion, knockout of EphA2 was confirmed by flow cytometry and the knockout of EphA4 was confirmed by Western blotting.

### Fusion assay

The virus-free cell-based fusion assay was performed as described previously described (42). Briefly, CHO-K1 cells grown to approximately 80% confluency in a 6-well plate and were transiently transfected with T7 luciferase reporter plasmid with a T7 promoter (1.5 μg) and the essential glycoproteins for EBV fusion gB (0.8 μg), EBV gH (0.5 μg), EBV gL (0.5 μg) or, for KSHV fusion, EBV gB (0.8μg), KSHV gH (0.5μg) KSHV gL (0.5μg) by using Lipofectamine 2000 transfection reagent (Invitrogen) in Opti-MEM (Gibco-life technology) as previously described (26). HEK293T cells or EphA2 single, EphA4 single, or double knockout (DKO) cells were transfected with T7 polymerase (1.5 μg) plus 1.5 μg pcDNA 3.1, EphA2 or EphA4 for the fusion assay. After 24 hours post-transfection, the transfected CHO-K1 cells were detached, counted, and mixed in a 1:1 ratio with target cells (HEK293T cells, 2.0 x 10^5^ per sample) into a 48-well plate in 0.5 mL Ham’s F-12 medium with 10% heat-inactivated FBS. 24 hours later, the cells were washed once with PBS and lysed with 50 μL of passive lysis buffer (Promega). Luciferase activity was quantified by transferring 20 μL of lysed cells to a 96-well plate and adding 50 μL of luciferase assay reagent (Promega). Luminescence was measured on a Perkin-Elmer Victor II plate reader. For the split GFP fusion assay, T7 polymerase was replaced with RLuc8 8-11 and T7 luciferase was replaced with RLuc8 1-7. The plasmids used were kindly provided by Gary Cohen and constructed by Matsuda and colleagues (43, 44). Images monitoring fusion were taken using an EVOS cell imaging system at 10X magnification.

### Cell surface expression

Surface expression of EphA2 was determined by flow cytometry analysis. 1x 10^6^ WT HEK293T cells, EphA2/EphA4 single, or double knockout cells were harvested and washed with PBS containing 1% bovine serum albumin (BSA) and incubated with 5 μL of PE-conjugated EphA2 antibody (Biolegend, SHM16) in 50 μL PBS containing 1% BSA for 30 minutes at 4 °C. Cells were then washed and diluted in 300 μl PBS containing 1% BSA. Data were acquired using a BD LSR Fortessa instrument and FlowJo software was used for analysis.

### KSHV infection

iSLK.219 cells (40) were cultured in DMEM supplemented with 10% fetal bovine serum (Tet-free FBS), 1% penicillin-streptomycin, 1 μg/ml puromycin, 250 μg/ml G418, and 1 mg/ml hygromycin B. KSHV producing cells, 2.0×10^5^ /well iSLK219 cells, were seeded in a 24-well plate in DMEM supplemented with 10% fetal bovine serum (FBS) and 1% penicillin-streptomycin. 24 hours later the cells were induced with 1 μg/ml doxycycline. After 4 days, the supernatant was collected and cells were pelleted at 1500 RPM for 10 minutes; the supernatant was aliquoted in 1 mL and then frozen at −80 °C or centrifuged at 13000 RPM for 30 minutes at 4 °C. The pellets were resuspended in 100 μL of 10% FBS DMEM. 7.5×10^4^ HEK293T cells/well were seeded on a 48-well plate and infected on the second day with 100 μL KSHV in 10% FBS DMEM. The percentage of infected cells was determined by flow cytometry or microscopic imaging.

### Western blotting

Expression of EphA4 was examined by Western blotting analysis. WT HEK293T cells and EphA2/EphA4 single or double knockout cells in 6 well plates were collected and resuspended in 50 ul PBS, then mixed with 50 μl 2X SDS loading buffer (60 mM Tris-Cl pH 6.8, 0.2% SDS, 25% glycerol, 0.01% bromophenol blue). Samples were boiled for 3 minutes and loaded onto a BioRad 4-20% mini PROTEAN TGX gel for western blotting. After electrophoresis, proteins were transferred to nitrocellulose membranes (Schleicher & Schuell, Keene, NH). The blots were blocked with 5% nonfat dry milk in PBS buffer (20 mM Tris-HCl, pH 7.6, 137 mM NaCl,) for 2 hours at room temperature (RT). The blots were washed with PBS and incubated with primary antibodies (anti-EBV gH/gL (a rabbit polyclonal antiserum;1:200) (45) and anti-His tag antibody (OB05, Calbiochem, 1:1000) for His-tagged KSHV gL) overnight at 4°C. Anti-rabbit IRDye800 or anti-mouse IRDye680 secondary antibodies (LI-COR bioscience, Lincoln, NE) were added to the membranes at a dilution ratio of 1:10,000 and incubated for 1 hour at RT. For detection of EphA2-Fc and EphA4-Fc, the membrane with transferred proteins was incubated with anti-human IgG (H&L) (HRP) (ab6759; Abcam, 1:1000) against the Fc region. The membrane was then incubated with 1mL SuperSignal chemiluminescent substrate (Thermo Fisher Scientific) prior to imaging. Protein bands on the membrane were visualized with the Odyssey Fc Western blotting imager using Image studio version 2.0 (LI-COR bioscience, Lincoln, NE).

### Cell enzyme-linked immunosorbent assay (CELISA)

The EphA2-Fc and EphA4-Fc bound to the transfected cells was determined by CELISA (cell enzyme linker immunosorbent assay). CHO-K1 cells were transiently transfected with control plasmid pcDNA 3.1, EBV gH/gL, and KSHV gH/gL. Soluble EphA2-Fc and EphA4-Fc were prepared by transfecting the CHO-K1 cells with EphA2-Fc and EphA4-Fc constructs and used to overlay epithelial cells (5×10^4^ cells/well) in 96-well plates in triplicate. After incubation for 2 hours at 4°C, the cells were incubated with anti-human IgG (H&L) (HRP) (ab6759; Abcam, 1:1000) against the Fc region for 30 minutes and fixed with 2% formaldehyde and 0.2% glutaraldehyde in PBS for 15 minutes followed by three PBS washes. TMB one component HRP microwell substrate was added and the amount of bound EphA2-Fc and EphA4-Fc was determined by measuring absorbance at 380 nm with Perkin-Elmer Victor plate reader. Binding activity was standardized in comparison to EphA2-Fc binding to EBV gB which was set to 100%.

### Statistical analysis

Data were collected from three independent experiments. Statistical differences between multiple groups were determined by one-way ANOVA with post-hoc Tukey’s multiple comparison test. Two-group comparisons were analyzed by the two-tailed unpaired Student’s *t* test. P < 0.05 denotes the presence of a statistically significant difference. Data are expressed as mean ± SE. The analysis was performed using GraphPad Prism, version 6.0c for Mac (GraphPad Software, San Diego, California, USA). Flow cytometry histograms and microscopy images are representative of at least two independent experiments.

### Data availability

The data that support the findings of this study are available within this article and its supplementary information files or upon requesting the relevant information from the corresponding author.

## Acknowledgements

We appreciate the help and advice from members of the Longnecker and Jardetzky laboratories, especially Nanette Susmarski. This research was supported by AI076183 (R.L. and T.J.), AI137267 (R.L. and T.J.) from the National Institute of Allergy and Infectious Diseases, IRG-15-173-21 (J.C.) American Cancer Society, and the Third Coast Center for AIDS Research pilot award (J.C.). We thank Gary Cohen for kindly providing the RLuc8 plasmids that were originally constructed by Z. Matsuda.

## Contributions

J.C. and R.L. designed the overall study with input from the co-authors. J.C. performed the key experiments. X.Z. performed the RNA seq analysis, aided in the design of the sgRNA, EphA2-Fc, and EphA4-Fc constructs, and with statistical analysis. S.S. helped with cell cultures and fusion assays and the generation of EphA2/A4 chimeras. J.C. and R.L. wrote the manuscript. S.S., X.Z., and T.J. contributed expertise and helped write the paper. All authors analyzed the results, read, and approved the manuscript for submission.

